# PEPOP: new approaches to mimic non-continous epitopes

**DOI:** 10.1101/435966

**Authors:** Vincent Demolombe, Alexandre de Brevern, Liza Felicori, Christophe NGuyen, Ricardo Andrez Machado de Avila, Lionel Valera, Bénédicte Jardin-Watelet, Géraldine Lavigne, Aurélien Lebreton, Franck Molina, Violaine Moreau

## Abstract

Bioinformatics methods are helpful to identify new molecules for diagnostic or therapeutic applications. For example, the use of peptides capable of mimicking binding sites has several benefits as replacing a protein difficult to produce, or toxic. Using peptides is less expensive. Peptides are easier to manipulate, and can be used as drugs. Continuous epitope predicted by bioinformatics tools are commonly used and these sequential epitopes are used as such in further experiments. Numerous discontinuous epitope predictors have been developed but only two bioinformatics tools proposed so far to predict peptide sequences: Superficial and PEPOP. PEPOP can generate series of peptide sequences that can replace continuous or discontinuous epitopes in their interaction with their cognate antibody. We have developed an improved version of PEPOP dedicated to answer to the experimentalists’ need for a tool able to handle proteins and to turn them into peptides. The PEPOP web site has been reorganized by peptide prediction category and is therefore better formulated to experimental designs. Since the first version of PEPOP, 32 new methods of peptide design were developed. In total, PEPOP proposes 35 methods in which 34 deal specifically with discontinuous epitopes, the most represented epitope type in nature.

We present the user-friendly, well-structured web-site of PEPOP and its validation through the use of predicted immunogenic or antigenic peptides mimicking discontinuous epitopes in different experimental ways. PEPOP proposes 35 methods of peptide design to guide experimentalists in using peptides potentially capable of replacing the cognate protein in its interaction with an Ab.

## Introduction

The antigen-antibody (Ag-Ab) interaction is the basis of the immune system, and the antibody a valuable tool in various biomedical applications, including diagnosis and therapy research [1,2]. The Ab plays a key role in two phenomena: immunogenicity and antigenicity. Immunogenicity is the ability of a molecule to induce an immune response in the host, yielding Abs. Antigenicity is the ability of a molecule to bind specifically to an Ab. Antibodies are known to exhibit highly specific binding, however, so off-target binding can occur [3]. The paratope of the Ab interacts with the epitope of the protein Ag. An epitope can be continuous or discontinuous, linear or conformational [4–6]. A continuous, linear, or sequential, epitope is a fragment of the protein sequence. A discontinuous epitope is composed of several small fragments that are scattered in the protein sequence, but are close when the protein is structured. A conformational epitope has to be correctly structured to be recognized by the Ab and is often discontinuous although it can be continuous, for example, in the case of a constraint mimotope.

Epitope prediction tools have been developed for two major reasons [7,8]. First, to identify in the protein fragments hoped to be more efficient and specific than the rest of the protein in eliciting anti-protein Abs by immunization in a host. Second, to identify epitopes recognized by an existing Ab. These tools hope to overcome the difficulties in experimentally mapping epitopes on proteins [9,10] as the most accurate method is the 3D structural identification of the Ag-Ab complex by X-ray crystallography, which is a time-consuming and laboring procedure.

The first epitope prediction tools predicted continuous epitopes from the protein sequence using propensity scales based on different physico-chemical properties [11] such as hydrophilicity [12], flexibility [13], β-turns [14], surface accessibility (19), or antigenicity [16]. Despite improvement attempts in the methodology [17,18], among them the combination of properties [19], Blythe & Flower showed that the predictions are not better than chance [20]. It was supposed that because most of the epitopes are discontinuous [21,22], the tools did not sufficiently take into account this criterion. The epitope prediction tools should consider structural information and target the identification of discontinuous epitopes. It is only rather belatedly that researchers take an interest in considering the 3D structure of the protein [23–25]. New epitope prediction tools are regularly developed [26–29].

However, theoretically, a tool cannot predict an epitope. An epitope only exists thanks to the existence of the Ab recognizing it. Hence, a protein is potentially composed of a mosaic of overlapping epitopes [30,31]. It is therefore theoretically possible to generate Abs against any region of the protein surface. Thus, each region on the protein can be a potential epitope.

Important research developments in this field do not concern real “ab initio” epitope prediction tools but fast and efficient methods dedicated to the complex task of dealing with discontinuous epitopes (either in helping to map them or in proposing immunogenic peptide sequences). These new bioinformatics methods could help in dealing with the discovering of new molecules, such as biomarkers or therapeutics, resulting from the high-throughput technologies like proteomics [32,33]. They could provide solutions to characterize these new molecules by developing probes to capture them, by mapping epitopes, identifying interaction sites, finding peptide surrogates, etc. Despite the interest in using prediction tools, in the end, the experimentalist will use peptides, either for immunization or to replace the protein in the interaction with the Ab [34].

Compared to continuous epitopes which are synthesized as such, prediction of peptides mimicking discontinuous epitopes is more complicated as a correct combination of the elements composing the epitope has to be found to build the peptide (see supplementary data 1).

Moreover, it is known that the recognition of the Ab can be very sensitive to the sequence: only one mutation can alter the interaction [35]. Thus, using the relevant sequence is crucial. Only two bioinformatics tools proposed so far to predict peptide sequences using 3D information: Superficial [36] and PEPOP [37]. Superficial predicts continuous and discontinuous peptides representing a potential epitope. The tool determines accessible protein fragments in a defined region on the protein and gathers them in a peptide, adding residues to link the fragments between them. PEPOP is an antigenic and immunogenic peptide prediction tool. The first version of PEPOP proposed three different methods to design peptides and we showed that they can be used to generate anti-protein Abs [37] or to map epitopes [38]. In our new research, we focused on novel methods that predict peptides representative of discontinuous epitopes and we benchmarked them [39]. In this article, we present different studies showing how peptides can be used to mimic discontinuous epitopes using the new web site, PEPOP version 2.0. Peptides predicted by PEPOP have been used as immunogens to prepare anti-protein antibodies using one peptide targeting one specific region. They have also been used in pairs to target two distinct regions on the protein, allowing the capture of the antigen. Peptides predicted by PEPOP have then been used as antigens either to experimentally map an epitope or to find an inhibitor of an Ab-Ag interaction. We showed the interest of using peptides that can represent the cognate protein. The ensemble of these improvements was implemented in the improved web-site. PEPOP v2.0 is available at http://pepop.sys2diag.cnrs.fr/.

## Results

### Description of PEPOP

PEPOP [37] is an algorithm dedicated to the prediction of peptides able to replace a protein in its interaction with an Ab. To predict a peptide in PEPOP, the first step is to determine the surface accessible amino acids (aa) and continuous segments. The second step is to define an area which can be a potential epitope. These areas in PEPOP can be either clusters of segments clustered according to their spatial distances or patches of 10Å, 15Å, and varying radii. The last step in predicting a peptide is to assemble the segments or the aa from an area in an arrangement so that the Ab will recognize it. Thirty-four methods based on different algorithms combine aa or segments to propose linear peptides mimicking discontinuous epitopes (for more details see [39]).

PEPOP is available in an improved new version of the web site (Figure 1). The web interface is composed of 3 sections that can correspond to different ways to use PEPOP in experimental projects. Below are four examples using PEPOP to predict peptides and use them in experiments. Each user is free to imagine other ways to use these “discontinuous” peptides.

**Figure 1.**
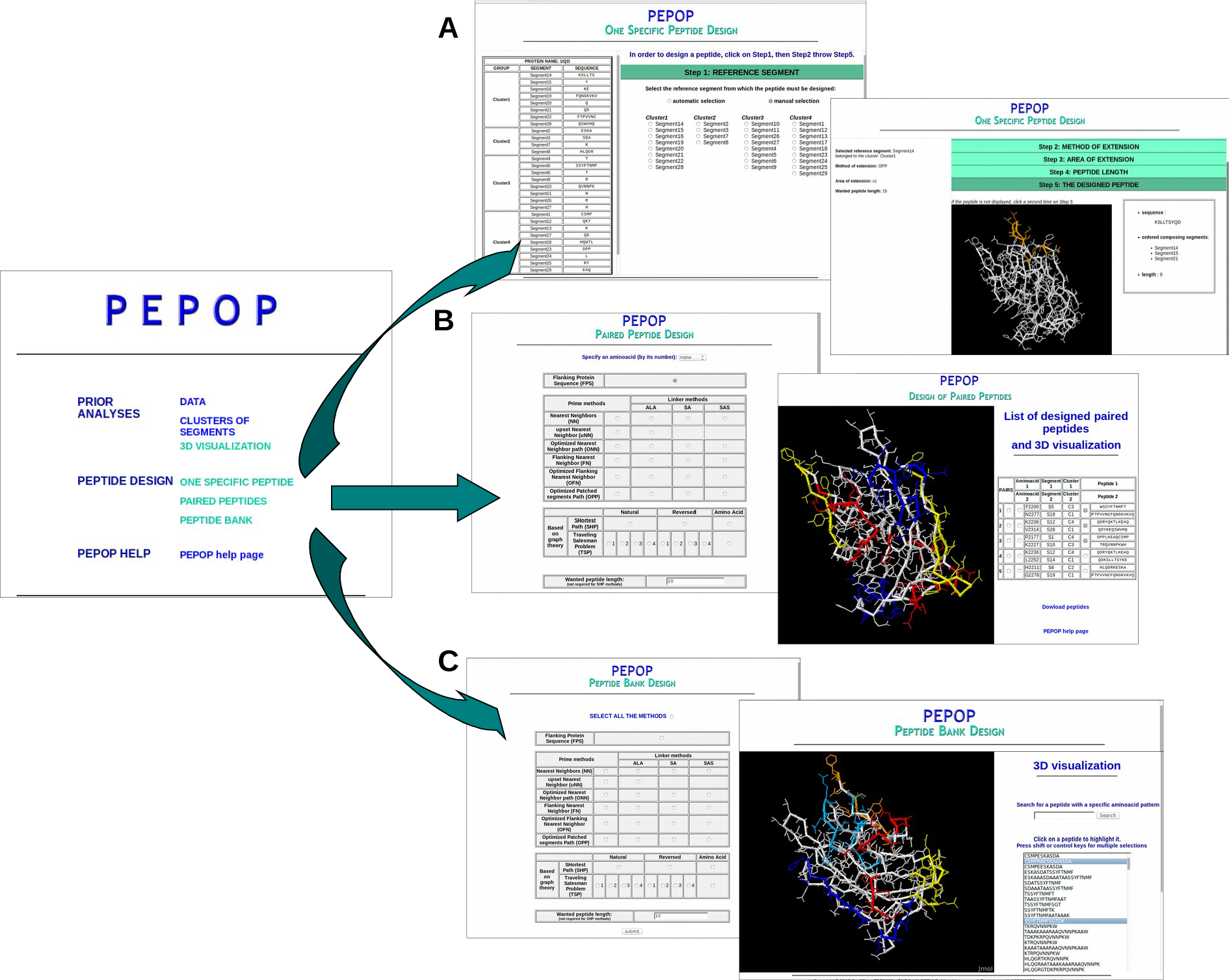
PEPOP web-site. The first result page of PEPOP, after the user gives the 3D structure of the protein, proposes 3 different ways to design peptides. **A**. The ‘One Specific Peptide Design’ predicts one peptide at a time through 5 steps where the user has to select the reference segment (first insert), the method of extension, the area of extension and the peptide length; the fifth step (second insert) gives the peptide sequence and displays it on the 3D structure of the protein. **B**. To design peptides in the ‘Paired Peptide Design’ section, the user selects the method of extension, the peptide length and eventually the aa from which the first pair has to be determined (first insert); the 5 peptide pairs are summarized in one side of the browser and displayed on the 3D structure of the protein on the other side of the browser. **C**. In the ‘Peptide Bank Design’, the user has to select the method(s) and the peptide length (first insert); all the predicted peptides can be displayed on the 3D structure of the protein (second insert).

The sections ‘One Specific Peptide Design’ and ‘Paired Peptide Design’ are dedicated to the prediction of peptides that will be used to generate anti-protein Abs. The ‘Peptide Bank Design’ section of the PEPOP web site is dedicated to the design of peptides that will be used for their antigenic properties. For this section, two types of experiments have been illustrated: the mapping of discontinuous epitopes and the identification of inhibitor peptides.

### Designing peptides to generate anti-protein Abs

The ‘One Specific Peptide Design’ section of the PEPOP web site is dedicated to the prediction of one peptide at a time. This section already existed in the previous version of PEPOP but was updated and enriched with new methods. This section allows defining only a small number of peptides. The peptide is progressively built through 4 steps: the reference segment, the method of extension, the area of extension and the peptide length. At each step, a choice is selected by default so that at the end the peptide can be built automatically. Else, the user may control the choices and the parameters (the 5 physicochemical and structural criteria: hyphobicity, accessibility, segment length, β-turn content, WRYP content) at any step. We prepared anti-protein Abs by designing a peptide from the 3D structure of the LMW (low molecular weight) form of adiponectin (PDB code: 1C3H) and using it to immunize mice. The peptide KYGDGDHNGLYADVETR has been predicted by the OFN method and gathered 4 segments: sequentially, segment 70 (K), segment 80 (YGDGDHNGLYAD), segment 81 (V), and segment 58 (ETR). We chose this method, new in this version of PEPOP, because we think it could be important to keep the reference segment in a central position in the peptide to be more easily recognized by the antibody. Figure 2 shows a dose-dependant reactivity of the Abs generated using this “discontinuous” peptide for the adiponectin. This result showed that PEPOP 2.0 sucessfully designs a peptide able to generate antibodies targeting a discontinuous epitope on the cognate antigen.

**Figure 2.**
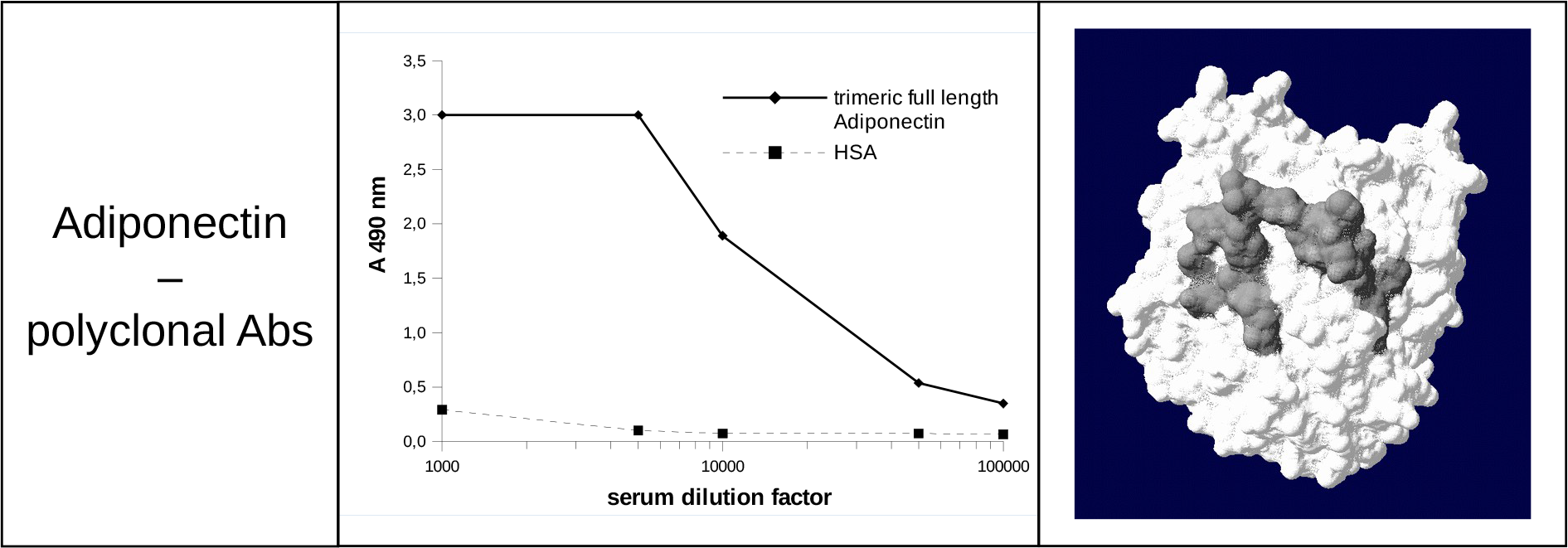
Reactivity of mouse immune serum raised to discontinuous adiponectin peptide against trimeric full length adiponectin (LMW adiponectin) and a control protein (HSA). The peptide is displayed on the surface of the protein.

### Designing peptides to generate Abs capturing the cognate protein

The ‘Paired Peptides Design’ section is new in this improved version of PEPOP. It is dedicated to the prediction of pairs of peptides. The goal is to target specific and distinct regions on the protein: the predicted peptides can then be used to prepare Abs that should be able to capture the cognate protein. The principle is to select two candidate peptides that are structurally appropriately separated in the 3D model. PEPOP proposes up to 5 pairs of distinct peptides. The peptides are designed by computing the most distant pairs of surface accessible aa and the two orthogonal most distant pairs in order to give the best chance to the generated Abs to capture the Ag without steric hindrance. Two more pairs are proposed as an alternative in the event that a targeted region is too close to the first one. This would lead to steric hindrance for the Abs generated. The user can orientate the design by indicating the position of one of the two aa of the first pair. The other pairs will be designed consequently. Figure 3 shows the example of the three first paired peptides on the A2 domain of FVIII. The six peptides are in distinct and opposite (two by two) regions of the protein. The recognition of the protein by the Abs generated by such peptides should not be disturbed by steric hindrance. The Abs should capture the protein two by two. This section of PEPOP can be a useful tool for the characterization of the proteins after a process of high throughput selection or for the development of a kit for diagnosis.

**Figure 3.**
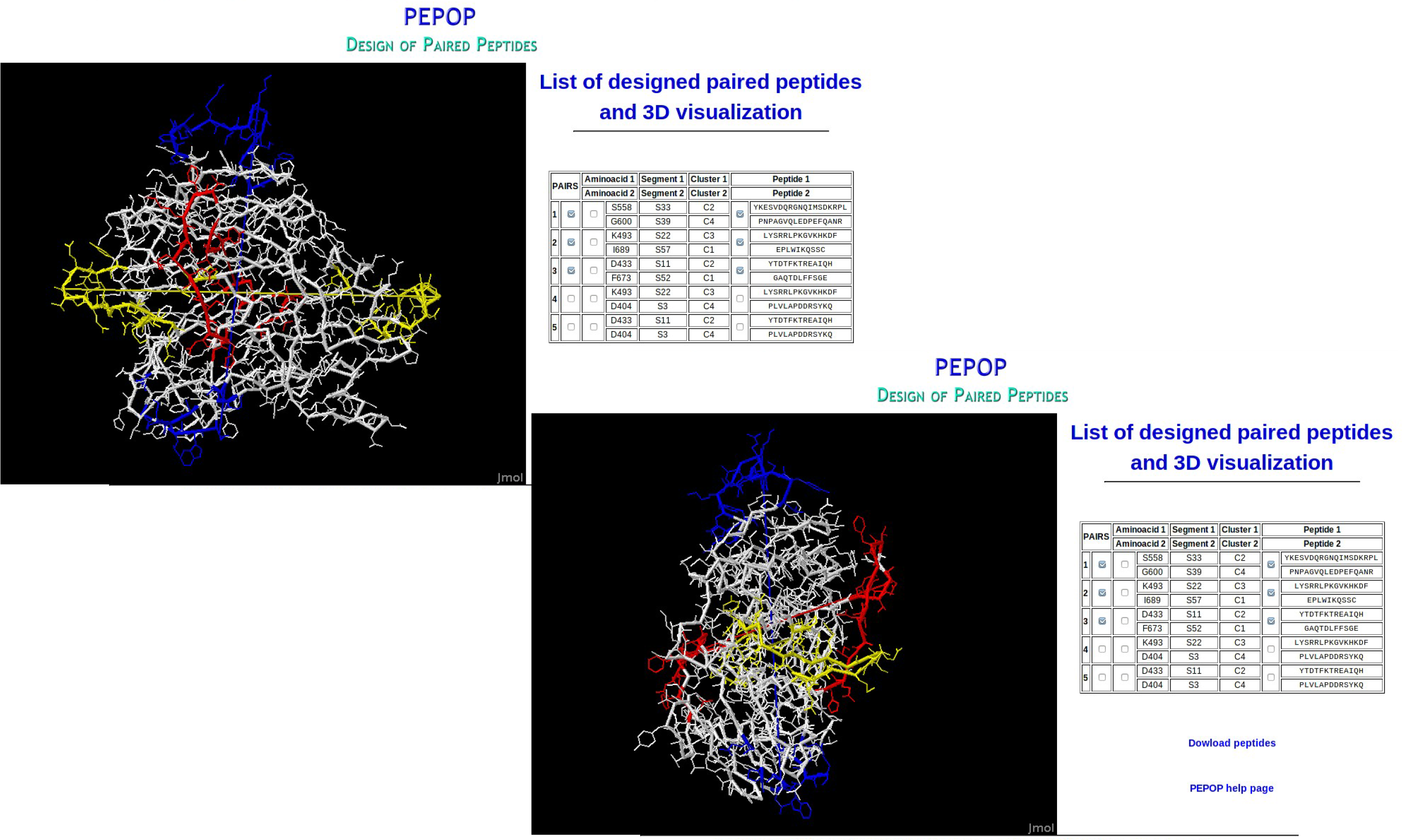
Example of paired predicted peptides on the A2 domain of FVIII. Paired peptides have been predicted from two distinct regions on the A2 domain of FVIII. The 6 peptides are in distinct and opposite (two by two) regions of the protein. The first paired peptides is in yellow, the second in blue and the third in red. The two 3D structure view are orthogonal.

We showed how PEPOP can propose peptides to use in immunogenic experiences. The designed peptides can also be used for their antigenic properties.

### Designing peptides to map discontinuous epitopes

The new section ‘Peptide Bank Design’ has been thought to propose an alternative to the existing heavy methods used to map discontinuous epitopes. The goal is a mixture of experiments used to map continuous (high-throughput peptide synthesis, e.g. SPOT technology [40,41]) and discontinuous epitopes (e.g. phage-display). As all the epitope information is already contained on the protein, we think experimental design is best suited by only testing the most numerous possible peptides, as in phage-display experiment. We drastically reduced the peptide space search by using protein information and methods carefully considered to address antigenic properties. The virtual peptide sequence bank is constructed thanks to a flexible web interface where the user has to choose the methods of extension and the peptide length (set to 10 aa length by default). Each method predicts all the possible peptides. For example, in the case of the prime, ALA, SA, and SAS methods, all the segments determined by PEPOP are individually selected as the reference segment. Thus, the method predicts as many peptides as segments. In this way, the entire surface of the protein is explored. Moreover, using several methods allows testing different arrangements of the same segments in peptides. Indeed, as we do not really know what governs the antigenic rules, we do not really know how some peptide characteristics, such as the peptide conformation, the aa position, the aa spacing, or the aa order influence the interaction with the Ab. The predicted peptides can be visualized on the 3D structure of the protein one or several at a time.

Using this methodology we map discontinuous epitopes either recognized by a pAbs on Amm8 [38] or recognized by mAbs on AaH II [35] and GM-CSF [42]. Figure 4 shows three more studies mapping discontinuous epitopes on LiD1 recognized by LimAb7 mAb and GAD65 recognized by DPC mAb and Ab54 mAb. Using prime, ALA, and SA methods with a requested peptide length of 10 aa, 456, and 648 peptides were predicted from the 3D model of LiD1 [43] and the 3D structure of GAD65 (PDB code: 2OKK) respectively. Peptides shorter than 7 aa have been eliminated because it is considered that the peptide is too short to well mimic the discontinuous epitope. Peptides longer than 24 aa have been eliminated due to synthesis performance limitations. Peptides have been synthesized using the SPOT method and their immune reactivities were tested with their respective mAb. In the case of LiD1, only one peptide has been recognized: it is displayed on the 3D structure of the protein. For GAD epitopes, several peptides have been identified. However, the control experiment with only anti-Fc pAbs reveals the reactivity of several peptides. By subtracting them, two specific spots appear that are only recognized by the mAb. According to the mAb, either DPC or Ab54, the two spots are different. The peptides representative of discontinuous epitopes are displayed on the 3D structure of GAD65.

**Figure 4.**
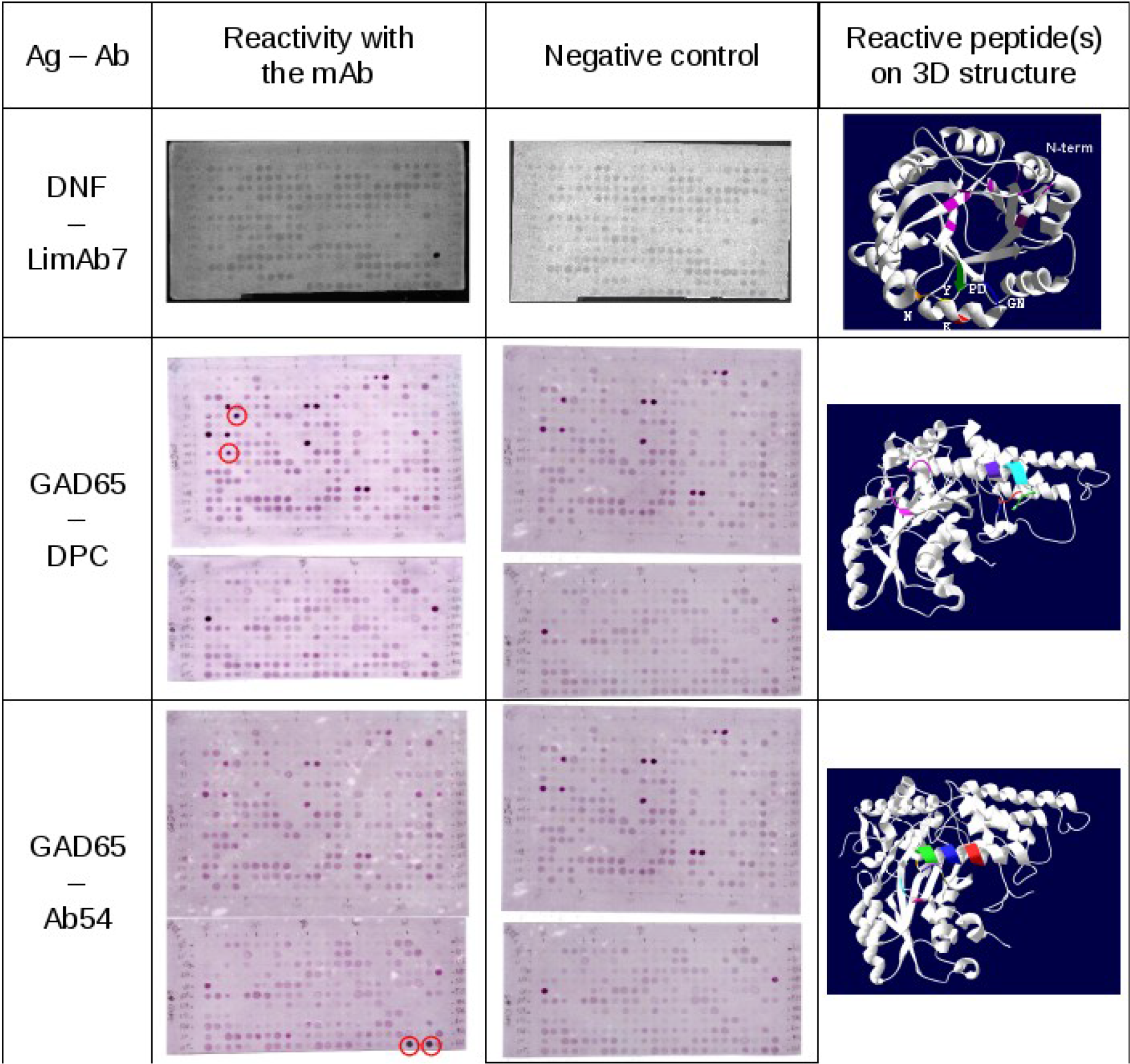
Reactivity of monoclonal antibodies, LimAb7, DPC and GAD65 with “discontinuous” peptides predicted from the 3D structure of respectively LiD1 and GAD65. The peptides have been prepared by the Spot technology. The reactivity was controlled with anti-Fc pAbs alone. The reactive peptides with the mAb are displayed on the 3D structure of the corresponding protein.

### Designing peptides to identify inhibitor peptides

Another way to use the ‘Peptide Bank Design’ section of the PEPOP web site is to test the antigenicity of the predicted peptides synthetized in soluble form with Abs in order to select peptides that could replace the cognate protein. Prediction of epitopes could have potential clinical implications in hemophilia A (HA), an inherited bleeding disorder. Indeed, severe HA is defined by an undetectable level of coagulation factor (F) VIII. The treatment of HA is based on regular intravenous infusions of FVIII and, to date, the main complication (up to 30 % of severe HA patients) of this treatment is the development of inhibitory anti-FVIII antibodies. The development of this immune response dramatically impacts the care of HA patients, and a fine epitope mapping could be helpful for a better understanding of the physiopathology and the treatment of such complications. As anti-FVIII Abs are mainly directed against C2 and A2 domain of FVIII, we predicted peptides mimicking discontinuous epitopes of these domains [44,45]. For example, we synthesised 33 synthetic peptides potentially representative of discontinuous epitopes on the C2 domain of coagulation FVIII, using the OPP method of PEPOP [44]. Only one method has been selected in the ‘Peptide Bank Design’ section. Indeed, as the experiments are relatively costly (in time and money) and need a large amount of plasma, all the peptides from the methods cannot be tested and a limited number of peptides needed to be selected. One solution is to select only one method. W chose this method because the reference segment is central in the patch, it contains no aa linker which could interfere with the Ab binding and the search of the path between the segments is optimized. In this way, the peptides together still allow exploring the entire surface of the protein. Using an inhibition assay based on the x-MAP technology, we evaluated their ability to block the binding to the C2 domain of anti-C2 domain Abs from plasma samples. Figure 5 shows one of the reactive peptides blocking the Ab binding in a dose-dependent manner. The peptides inhibits the interaction between the C2 domain of FVIII and the Abs by around 30%. The same protocol with another PEPOP method, TSPaa, was used to predict peptides mimicking discontinuous epitopes of the A2 domain of FVIII. For more details, see [45]. So, we show that it is possible to find at least one peptide in a series predicted by PEPOP that inhibits an Ab-Ag interaction.

**Figure 5.**
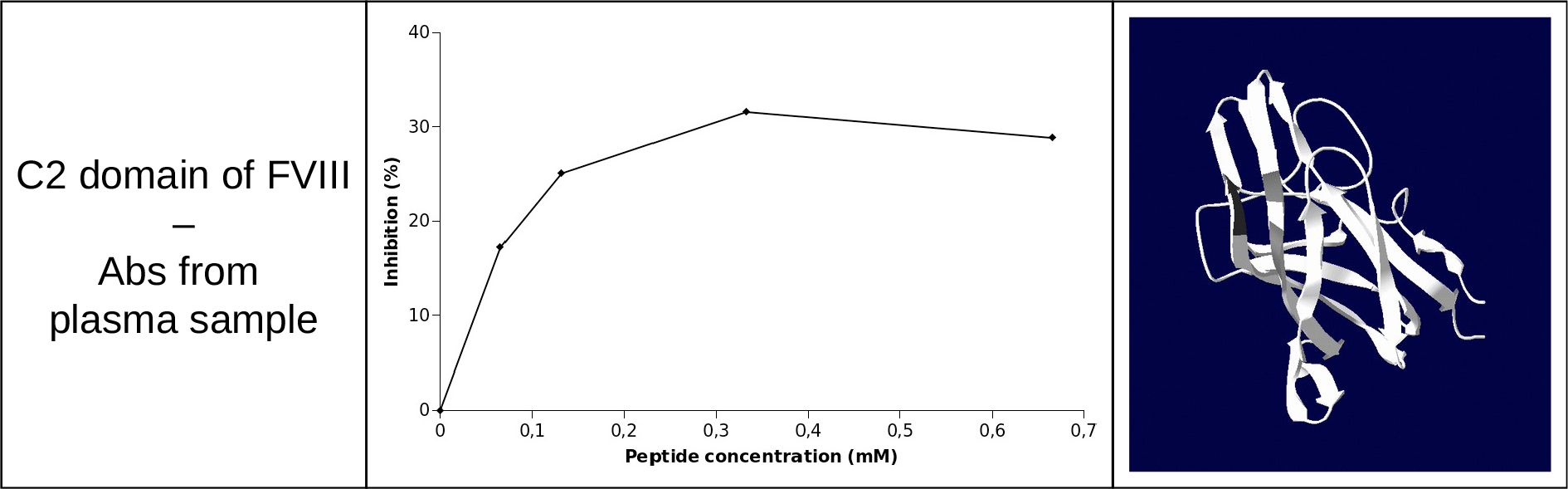
Inhibition obtained with different amounts of a peptide representative of the C2 domain of FVIII in x-MAP inhibition assays using plasma sample.

At any section of the PEPOP web site, the location of the predicted peptides can be displayed on the 3D structure of the protein.

## Discussion

By presenting the improved version of the PEPOP web-site, we showed the ways to use predicted peptides expected to mimic discontinuous epitopes. The most often use of the peptides is the generation of anti-protein Abs. One of the two great novelties of PEPOP is the use of peptides by pair so as to target distinct regions on the surface of the protein and generate Abs that should be able to capture the protein. This can be a useful tool, for example, in the characterization of biomarkers after the process of discovery in high-throughput selection. It could lead notably to the development of diagnosis kits. The other novel feature of PEPOP v2.0 is the ‘Peptide Bank Design’ section of the web-site. Because we predict from the native antigen, we showed that only a limited number of peptides (compared to the diversity generated in phage-display method) is necessary to map discontinuous epitopes. After synthesis, the functionality of the peptides exploring all the surface of the protein could be assessed in a convenient high-throughput recognition assay, such as the SPOT method [38] or other technologies [46]. If the correct sequence is present in the bank, the Ab should recognize it which identifies the epitope region on the protein. Then, a set of peptides around the space of the epitope region identified can be tested in further experiments to hone more precisely the epitope or to select a functional peptide. The final feature we tested is the search for an inhibitor. We synthesized, in soluble form, a restricted list of peptides and tested their capacity to inhibit the interaction between the protein and Abs. We showed that it is possible to select peptides able to replace discontinuous epitopes in an Ag-Ab interaction.

Two opposing views exist about epitopes. The first view considers that a protein is constituted by a mosaïc of potential epitopes [30]. The other point-of-view considers that proteins have only a few epitopes preferentially recognized by the immune system [47]. In view of these two hypothesis, it is not surprising that Blythe and Flower found that the continuous epitope prediction tools are not better than chance [20] and that the discontinuous epitope prediction tools showed weak performances [48]. In the first hypothesis, a tool cannot find any region emerging from the others since it is possible to produce Abs targeting any surface of the protein. In the other hypothesis, it would likely be logistically impossible for a tool to well predict when the learning data are a mix of a variety of different epitopes (immunogen, epitopes generated from peptides, truncated protein, cross-reacting molecules). The properties of Ag-Ab complexes have largely been analyzed but without distinguishing their types and origins. To know whether it is really possible to predict *ab initio* epitopes, the existence of immunodominant regions should be proved or refuted, for example with systematic studies by categorizing Ag-Ab complexes. Perhaps, we will discover that it is an intermediary or both of the two hypotheses: the immune system could preferentially target few specific regions on the protein (would it be just a question of surface accessibility?) but it still is possible to produce Abs targeting any regions [49]. Whatever the reality, in the present state of knowledge, the only way to predict an epitope is to take into account the Ab. One epitope only exists with the Ab that recognized it [50].

Predictings an epitope beings by proposing a region on the protein, i.e. a set of aa. Peptide prediction tools have to determine the sequence from this set by determining an arrangement, a disposition, a path between the aa. This can be very difficult. More elements have to be combined, and as the problem becomes more complex, it becomes rapidly unsolvable. This is an NP-complex problem relying on combinatorial mathematics. Solutions have to be found because it is impossible to edit all the possibilities.

Moreover, although the Ag-Ab interactions have been deeply studied [51–54], the mimicking of a discontinuous epitope by a linear peptide still is a challenging task [55]. Other parameters than those found in protein-protein interface studies [56–58] have to be taken into account. Should the peptide adopt the same conformation as in the protein so as the Ab can recognize it? Would the peptide be in the same conformation in the protein context? Chen et al. [53] showed that the conformations of the peptides compared to those of the corresponding regions on the proteins when complexed with the Ab have considerable differences. It should be even more difficult because the structure of an epitope when it is complexed with the mAb tends to differ from the structure before the mutual adaptation process [59]. Should the aa be spaced out as in the protein so that they are correctly laid out to allow the CDR loops of the Ab well facing them and interact with them? Or, is it sufficient for the key to be present in the peptide whatever their disposition? Actually, it is poorly known about the molecular mimetism. It should be very informative to carry out systematic studies in order to fully elucidate this phenomenon. In this way, PEPOP can be seen as a “test tube” to help to better understand molecular mimetism.

However, there is a real advantage in using mimicking peptides. Beyond avoiding the difficulties to obtain a pure preparation of the protein, reduction in cost, and increased ease in manipulation, even with polyclonal Abs the regions targeted on proteins are well known. The main advantage of using “discontinuous” peptides is that the final Abs should recognize the native well-structured protein antigen. Moreover, the same series of peptides can be probed by different Abs raised against the same target antigen, so as to disclose the cognate epitope of each.

However, the experimentalists have to carefully think through their experiments before designing peptides because, as van Regenmortel underlined at a workshop about current state and future directions for the epitope prediction field [60], the results can be different according to the experiment. For example, a peptide seen reactive in SPOT could be found not interacting in soluble form in ELISA. It may be due to the different conformation the peptide adopts according whether it is linked to a support or totally free in solution. It also may be due to the phenomenon of avidity in SPOT. Thus, if the experimentalist wants to map the epitope, he can carry on SPOT experiments or other high-throughput technologies. But, if he wants to use the reactive peptide in further experiments, he has to keep in mind that they could not react the same way. This is why it is recommended for experimentalist searching for an inhibitory peptide to select it by using technology that will present the peptides in its final format. Furthermore, the experimentalist also has to carefully choose the peptide design methods according to the objectives of the experiment. If the aim is to generate Abs, it would be better not to use linker methods in order to avoid that the Abs are directed against the linker aa, which could lead to Abs not cross-reacting with the protein. If the aim is to find an inhibitory peptide (experiments in soluble form), it is recommended to use peptide design methods that search for an optimized path (ONN, OFN, OPP, graph-based methods).

Assigning a function to each new protein structure resulting from high-throughput genomics experiments is a huge task. For example, the current techniques for epitope mapping are unfeasible on a genomic scale due to the high cost and effort needed. Reliable computational methods can assist by offering fast, scalable, and cost-effective approaches for identifying B-cell epitopes, focusing on experimental studies and improving our understanding of Ag-Ab interactions. PEPOP is a bioinformatic tool designing immunogenic or antigenic peptides representative of a given protein. It should facilitate experimentalists to handle proteins by turning them into peptides, smaller and easier to manipulate. Such a tool is in line with future goals in the area of the discovery of biomarkers by providing solutions to characterize these molecules or develop probes to capture them, leading to diagnosis and therapy applications. PEPOP will also be a useful tool to discover and study the rules governing molecular mimetism by testing the different approaches developed in the peptide design methods through systematic studies on antigenicity or immunogenicity. Furthermore, the tool is sufficiently flexible for allowing other problems to be addressed. For example, one can compare the peptides representing the surface of two proteins known to interact with the same mAb. Or, as PEPOP explores the surface of any protein, it can potentially be used to investigate any protein-protein interaction: the Ab would be replaced by another protein interacting with a targeted protein. Therefore, the protein-protein interaction site or an inhibitory peptide could be searched for the same way. This opens the door to an even greater world of possibilities in diagnosis or therapy applications.

## Experimental procedures

### Patch definitions

PEPOP uses three types of patches. The 10Å- and 15Å-radius patches gather the segments distant from the G point of a reference segment (selected by the user or by default) of a fixed distance, respectively 10Å and 15Å. In the third type, the patch gathers the aa distance from a reference amino acid (aa) of a distance varying from 15 to 20Å: the final radius corresponds to the one for which the number of aa collected is the average number of aa between radius 15, 16, 17, 18, 19 and 20Å.

### Prediction of paired peptides

The first step is to determine the distant aa. Then, the peptides are designed according to the method either by considering the aa (SHPaa and TSPaa methods) or the segment including it as the reference (starting point).

The first pair of distant aa is the two most distant surface accessible aa of the protein. To find the second pair of aa, an orthogonal plane to the first pair of aa is drawn. The two most distant aa around 5Å from this plane are searched for. A distance from the plane has to be tolerated else the plane could cross a zero aa threshold. The third pair is the two most distant aa included in the 10Å-thickness perpendicular bisector to the first and the second pairs of aa. A fourth and fifth pairs are proposed as an alternative to the second and third pair, respectively. The fourth pair of aa consists of one of the two aa of the second pair and the most distance of all the surface-accessible aa of the protein. The fifth pair of aa is the most distant pair where one is one of the two aa of the third pair and the other is found among all the surface-accessible aa of the protein.

### Web interface

PEPOP has been implemented on a virtualized Linux server kernel 2.6 running the Apache web server version 2.2.15. The tool has been implemented in object-oriented PHP and Javascript, and uses scripts and softwares developed in PERL, C, and C++. Segments, clusters, and peptides identified by PEPOP can be directly visualized on the 3D structure of the Ag thanks to jmol. PEPOP is available at http://pepop.sys2diag.cnrs.fr/.

## Experiments

### Synthesis of spot peptides

The peptide analogs were prepared by Spot synthesis [61] on a cellulose membrane, as previously described by Laune *et al.* [62]. Membranes were obtained from Proteigene. A Multipep robot (Intavis) was used for the coupling steps. Peptides were acetylated at the N-terminus. After the peptide sequences were assembled, the side-chain protecting groups were removed by trifluoroacetic acid treatment, but peptides remained attached on the membrane for ELISA-Spot experiments. Briefly, after an overnight saturation step with 3% BSA, the set of membrane bound peptides were probed by incubation with the mAb of interest. After 90 min incubation at room temperature, the membrane was washed and incubated for 1 h at room temperature with a peroxidase-conjugated anti-mouse or anti-human antibody (Sigma, diluted 1:3000). The spots were stained with enhanced chemiluminescent ECL detection kit (Amersham). The reactivity of each membrane was assessed in at least three independent experiments.

### Synthesis of the “discontinuous” adiponectin soluble peptide and the coagulation FVIII soluble peptides

The soluble peptides were synthesized on a Multipep Synthesizer using fluorenylmethyloxycarbonyl (Fmoc) acting as a protective group [63,64] with a HOBt-DIPC protocol. The C-terminal residues were first fixed to the solid phase support and NH2 extremity, and R groups are initially protected by Fmoc. After a basic deprotection of the NH2 extremity of the first fixed aa, the second protected aa was added and its carboxyl function activated to allow for the peptide linkage and extension of the peptides. The peptides were elongated after a succession of protection/deprotection steps until the addition of the last residue. Lateral chains were subsequently deprotected and the peptides released from the resin by trifluoroacetic acid treatment in the presence of the appropriate scavengers in order to generate amidated peptides. After synthesis, the peptides were lyophilized and the quality of the peptides was verified by high performance liquid chromatography and mass spectrometry.

### Immune response to the “discontinuous” adiponectin peptide

#### Mouse immunization

Eight-week-old Balb/C male mice were immunized by intraperitoneal injection of complete Freund adjuvant (first injection) and incomplete Freund adjuvant (following injections) containing KLH-conjugated discontinuous adiponectin peptide. After the 5th injection, blood was collected in order to characterize the immune response to full length adiponectin. The study was approved by the “direction départementale de la protection des populations” (B34-172-27 agreement).

#### Immune response characterization

Binding activity of the immune serum to recombinant adiponectin and irrelevant protein was evaluated by direct ELISA. The plates were coated overnight at 4°C with 1μg/ml of recombinant trimeric form of human adiponectin produced in HEK cells (BioVendor, #RD172091100) or purified Human Serum Albumin (HSA) (Sigma-Aldrich, A9511). After blocking with 1% milk in PBS, mouse serum was diluted from 1/1000 to 1/100 000 in PBS with 0.1% milk and 0.1% Tween and plates were incubated for 1h. After washing, the secondary antibody Peroxidase-AffiniPure Donkey Anti-Mouse IgG (Jackson Immuniresearch, 715-035-150) was incubated for 1h at 1/3000 in the same buffer followed by o-Phenylenediamine dihydrochloride (Sigma Aldrich, P8412). The absorbance was measured at 490 nm.

### Plasma reactivity with the discontinuous peptides from coagulation FVIII

The ability of peptides to inhibit the binding of anti-C2 Abs to the C2 domain was then evaluated in an original and homemade inhibition assay. Briefly, the domain of interest was immobilized on luminex beads. The plasma of hemophilia A (HA) patients containing anti-C2 Abs was incubated with a range of concentrations of peptides and thereafter incubated with beads. The dose-dependent inhibition of peptides was revealed with a fluorescent anti-human Ab, recognizing residual plasma Abs bound to the specific domain coated on beads if the predicted peptide mimicked one of the epitopes recognized by human anti-C2 Abs, the level of fluorescence decreased and the inhibition rate increased. This study was connected to Lapalud et al study [65] for which the protocol was approved by the Ethics Committee of Montpellier (France), and informed consent was obtained for all patients in accordance with the Declaration of Helsinki.

## Acknowledgements

We warmly thank Dr P. Lapalud and Dr B. Jardin for hers collaborations and D. Jean for his precious technical contribution. This work was supported by grants from the Ministry of Research (France); University Paris Diderot, Sorbonne Paris Cité (France); the National Institute for Blood Transfusion (INTS, France); the brazilian instituitions CAPES, FAPEMIG and CNPQ; the Institute for Health and Medical Research (INSERM, France); and a Labex GR-Ex grant (France) to Alexandre de Brevern. The Labex GR-Ex, reference ANR-11-LABX-0051, is funded by the program “Investissements d’avenir” of the French National Research Agency, reference ANR-11-IDEX-0005-02. We are grateful to A.-M. Madec and M. Powell for providing anti-GAD65 DPC and mAb54 monoclonal antibodies.

